# An ALS Assembly Modulator Signature in Peripheral Blood Mononuclear Cells: Implications for ALS Pathophysiology, Therapeutics, and Diagnostics

**DOI:** 10.1101/2024.03.28.587289

**Authors:** Shao Feng Yu, Maya Michon, Anuradha F. Lingappa, Kumar Paulvannan, Dennis Solas, Kim Staats, Justin Ichida, Debendranath Dey, Jeffrey Rosenfeld, Vishwanath R. Lingappa

## Abstract

Assembly modulators are a new class of allosteric site-targeted therapeutic small molecules, some of which are effective at restoring nuclear localization of TDP-43 in ALS cellular models, and which display efficacy in a variety of ALS animal models. These compounds have been shown to bind selectively to a small subset of protein disulfide isomerase (PDI), a protein implicated in ALS pathophysiology. The targeted subset of PDI is found within a novel, transient and energy-dependent multi-protein complex that includes other important members of the ALS interactome, such as TDP-43, RanGTPase, and selective autophagy receptor p62/SQSTM1. We demonstrate here that a similar multi-protein complex drug target is present in PBMCs as isolated by energy-dependent drug resin affinity chromatography (eDRAC) and characterized by mass spectrometry and by Western blot (WB). Signature alterations in the composition of the multi-protein complex in PBMCs from ALS patients compared to PBMCs from healthy individuals were identified by WB of eDRAC bound proteins, thereby extending earlier literature suggesting PBMC dysfunction in ALS. Changes in the PBMC drug target in ALS patients compared to healthy individuals include diminished p62/SQSTM1 and appearance of a 17kDa post-translationally modified form of RanGTPase. These changes are not readily apparent from analysis of whole cell extracts, as the individual protein components within the drug target multi-protein complex comprise only small percentages of the total of those component proteins in the extract. Furthermore, whole blood from ALS patients shows a distinctive degradation of total RanGTPase not observed in blood from healthy individuals. This degradation appears to be rescued by treatment of whole blood from ALS patients for 72 hrs with ALS-active assembly modulator small molecules. Our findings are consistent with the hypothesis that ALS is fundamentally a disorder of homeostasis that can be detected early, prior to disability, in blood by the methods described, and restored to the healthy state by assembly modulator drug treatment.

## Introduction

Amyotrophic Lateral Sclerosis (ALS) is a devastating neurodegenerative disease that strikes approximately 6,000 American and 150,000 people worldwide annually^1,2^. Approximately 10% of ALS is familial, with a large number of specific gene mutations implicated. The vast majority of ALS cases however, are sporadic, meaning that that there is no family history of the disease and therefore seems likely due to an environmental cause, perhaps with epigenetic influence^3^. Specific environmental factors have been implicated in causing ALS, but remain controversial and inconclusive^4^. While the pathophysiology of ALS is not well understood, aggregates of the protein TDP-43, independently implicated in the HIV life cycle^5^, are almost invariably found in the brains of patients with ALS and fronto-temporal dementia (FTD)^6^. Despite this commonality, the clinical presentation as well as progression of ALS, is highly variable^7^.

With regards to TDP-43 pathophysiology, controversy exists as to the triggering event. Some studies have implicated mislocalization of TDP-43 from the nucleus in healthy cells to the cytoplasm, which is observed to variable degrees in fibroblasts from many, but not all, cases of ALS^8^. An alternative view is that, in response to stress, TDP-43 forms aggregates in the cytoplasm where it is synthesized, prior to transport to the nucleus^9^. A further controversy exists as to whether the relevant TDP-43 aggregates in the cytoplasm are within, or outside of, stress granules^10^. While the TDP-43 aggregates have generally been viewed as a trigger of motor neuron death, it is possible that they are an epiphenomenon whose elimination *per se* may not necessarily be therapeutic, by analogy to the failed attempts to remedy Alzheimer’s Disease by elimination of Aβ aggregates^11^. Regardless of their natural history and role in the pathophysiology of ALS, the presence of TDP-43 aggregates are pathognomonic for the disease^12,13^.

Recently we presented our studies on the role of a newly appreciated dimension of gene expression, termed protein assembly, in the pathophysiology of diverse diseases including ALS^14,15,16^. These compounds were found by a novel method in which viruses were used as “trufflehounds” to reveal drug targets inaccessible by genomic or conventional proteomics^15,16^. We have identified several structurally unrelated small molecule chemotypes by virtue of their ability to modulate host machinery catalytic for viral capsid formation, that we term protein assembly modulators. These compounds are strikingly efficacious both for the specific viral disease by which they were identified, and also for a subset of non-viral diseases. Among the protein assembly modulators with activity against HIV, one prevents ALS-associated neurodegeneration and its correlates in various cellular and animal models of ALS^14^. A subseries of analogs shows diminished activity against HIV and improved activity against ALS. These compounds are broadly effective at reversing both TDP-43 mislocalization and stress-induced TDP-43 aggregation, in both familial and sporadic ALS and fronto-temporal dementia (FTD) cellular models. Efficacy has also been demonstrated in various transgenic animals expressing human ALS-causing mutations including in TDP-43, FUS, c9orf72, and SOD1, by a variety of assays^14^. In view of this breadth of efficacy, we hypothesized the drug target to be a step common to most, if not all, ALS and FTD. Advanced analogs of the ALS-selective active subseries of one of these chemotypes are close to completion of a target product profile (TPP), prior to initiation of investigational new drug (IND)-enabling studies in anticipation of human clinical trials in the near future.

The studies to be described here had several motivations. First, to explore the hypothesis that the biochemical pathways involved in ALS pathophysiology may involve layers of regulation yet to be discovered^17,18^, in which the novel multi-protein complex targets of these compounds might be involved. How those levels of control are integrated with one another, and are communicated from one organ system to others, remain largely unknown. It may be that the novel assembly modulator targets we have discovered are potential candidates for involvement in such regulation. By corollary, what happens to other feedback controls when one level of regulation fails, remains to be determined. The argument has been made based on gene knockout experiments that more ancient back up pathways may be activated when primary regulatory controls fail^19,20^. Having a drug that impinges on a biochemical pathway in a novel way is a powerful means of probing biological regulation. Much of this regulation depends on information that does not reside simply in the sequence of the genome^18,20,21^. Thus the relevance of the compounds we describe, with their unconventional multi-protein complex target (energy-dependent for formation, transient in existence, composed of miniscule subsets of the individual component proteins in the cell), and novel mechanism of action at allosteric sites^22,23^ ^24,25^. These tools may provide a new understanding of the molecular basis for homeostasis, and its dysregulation, as exemplified in ALS.

A second motivation for the present studies was to understand the meaning of the observed dysfunction of PBMCs in ALS^26,27^. These findings can be interpreted in very different ways. On the one hand, PBMC dysfunction might be due to ALS being a systemic disease whose most severe manifestation is motor neuron death, but which also manifests in less dramatic fashion in other organ systems, including in blood. Alternatively, the signature observed in PBMCs might reflect a role of blood, which sees essentially every cell in the body on a regular basis, spreading the message that there is a disorder affecting a particular cell type, in this case, motor neurons. Yet another possibility is that the disorder affecting one organ system (motor neurons) could have a “domino effect” manifesting in other organ systems (PBMCs), e.g. as a secondary consequence of feedback regulation intended to monitor and maintain homeostasis^17,28^. Prion-type spread observed for various diseases may be an example of such disordered communication^28–30^. To better understand these alternatives, we asked whether assembly modulator drugs might detect a version of the disordered drug target in PBMCs from ALS patients.

Finally, apart from intellectually interesting questions of feedback and mechanism of action^17,28,30^, there is a practical desire to advance these compounds to the clinic, given their remarkable therapeutic properties in cellular and animal models of ALS^31^. Having a readily accessible signature of ALS in PBMCs would facilitate ALS drug advancement in innumerable ways, most importantly by making possible early detection of ALS (prior to severe disability). Compounds designed to restore homeostasis would be expected to be most effective when administered early, before motor neuron death. Or put another way, treatments that restore homeostasis would be expected to be very different from those needed to regrow and replace cells after they have died. A signature in PBMCs would also provide a biomarker for easy assessment of therapeutic efficacy, and to determine when therapy could be stopped or might need to be restarted.

## Results

Blinded blood samples were obtained from ALS patients and healthy individuals, under an IRB-approved protocol. The PBMCs were isolated, and extracts prepared as described in methods, and analyzed for key proteins relevant to ALS. Differences observed in the protein pattern between PBMCs of healthy and ALS patients, by total protein silver stain were subtle (see for example, **Figure 1A**).

**Figure 1.**
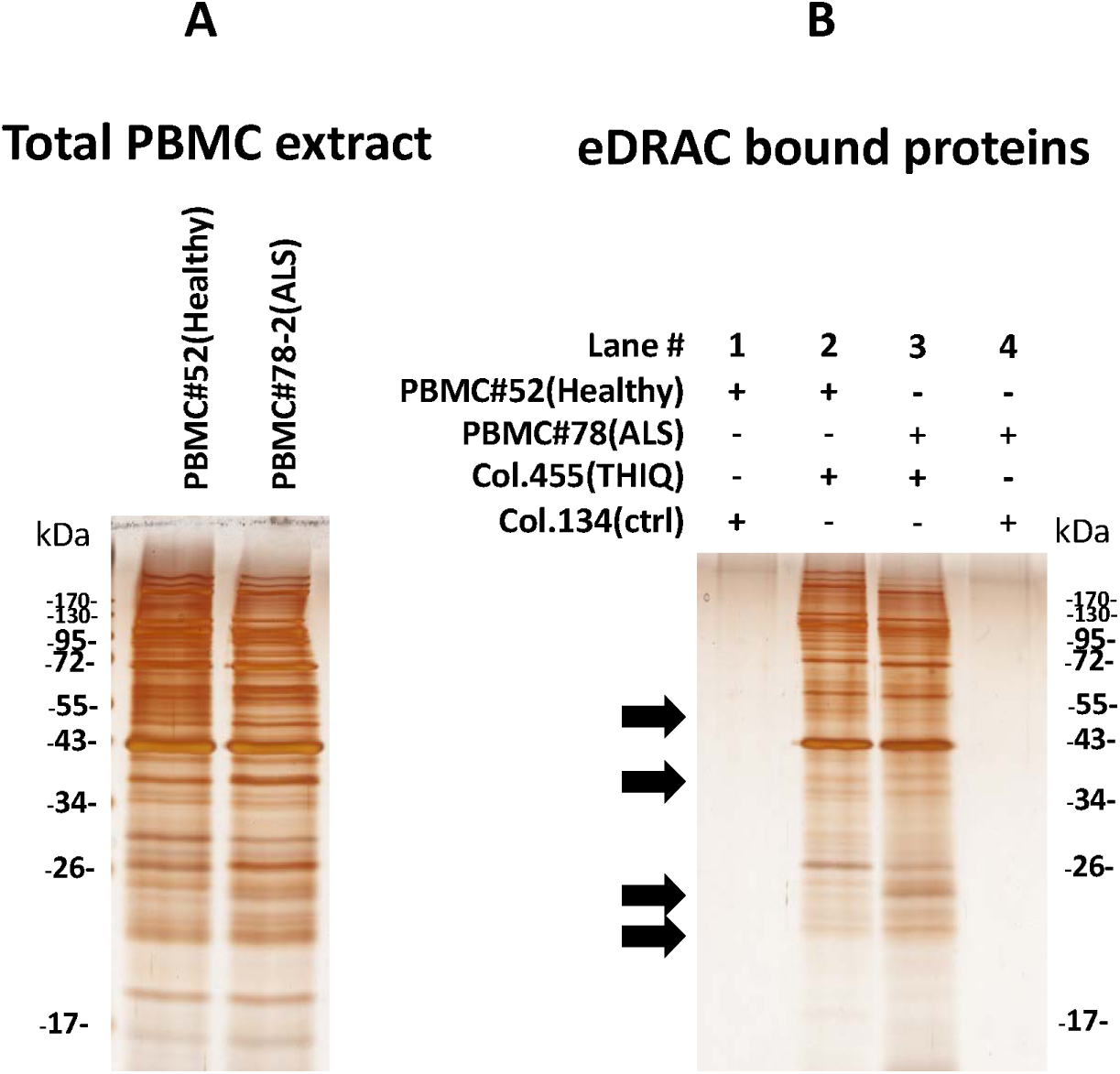
A. Silver stain of starting PBMC extract 10kxg/10min supernatant from a representative healthy individual (#52) and an ALS patient (#78-2). B. Silver stain of free drug eluate from ALS-active assembly modulator drug resin (column 455 THIQ) or control resin (column 134 lacking the drug ligand). Black arrows indicate positions at which healthy vs ALS patient samples show a suggestive difference in the free drug eluate protein pattern.

However when PBMC extracts were applied to the ALS-active drug resin described previously^31^ versus control resin (lacking the drug ligand), washed and bound material analyzed, a distinctive subset of proteins was observed, also largely shared between healthy individuals and ALS patients, but with a few possible differences (black arrows in **Figure 1B**).

Previous studies of assembly modulator small molecules^15,16^ have shown that the proteins comprising the multi-protein complex drug target represent single digit percent of the total of those proteins in the starting extract. As expected therefore, the depleted extracts differ very subtly from the starting material, as demonstrated for PBMC DRAC flow through here (see also **Supplemental Figure 1D**).

Mass spectrometry analysis of free drug eluates from PBMCs applied to the ALS-active drug resin (**Figure 2**) was carried out and compared to the pattern previous revealed from mouse brain (MoBr). A substantial proportion of tissue-specific proteins (~1/2 to 2/3 depending on the specific sample) were observed (see **Supplemental Figure 2**). Despite the high tissue specificity, a core set of 33 proteins shared between both PBMCs and brain were of interest. Of these, 26 were identified as members of the ALS protein interactome. Notably, this set included P4HB, a member of the PDI family, two known ALS-causing proteins (TUBA4A and VCP)^32^, along with proteins implicated in the KEGG ALS interactome^33^ (ATP5A1, HSPA5, TUBA1A, TUBA4A, and VCP). Furthermore, 24 of these 33 proteins exhibited high-confidence interactions with known ALS-causing or disease-modifying proteins^32^. The 26 shared ALS interactome proteins also included multiple members of several protein families and complexes: two ATP synthase subunits (ATP5A1 and ATP6V1A), six chaperonins containing T-complex components (CCT2, CCT3, CCT4, CCT5, CCT7, and CCT8), three thioredoxin domain proteins (P4HB, PRDX1, and PRDX2), and four members of the 14-3-3 family (YWHAB, YWHAE, YWHAH, and YWHAQ) (**Figure 2A**).

**Figure 2.**
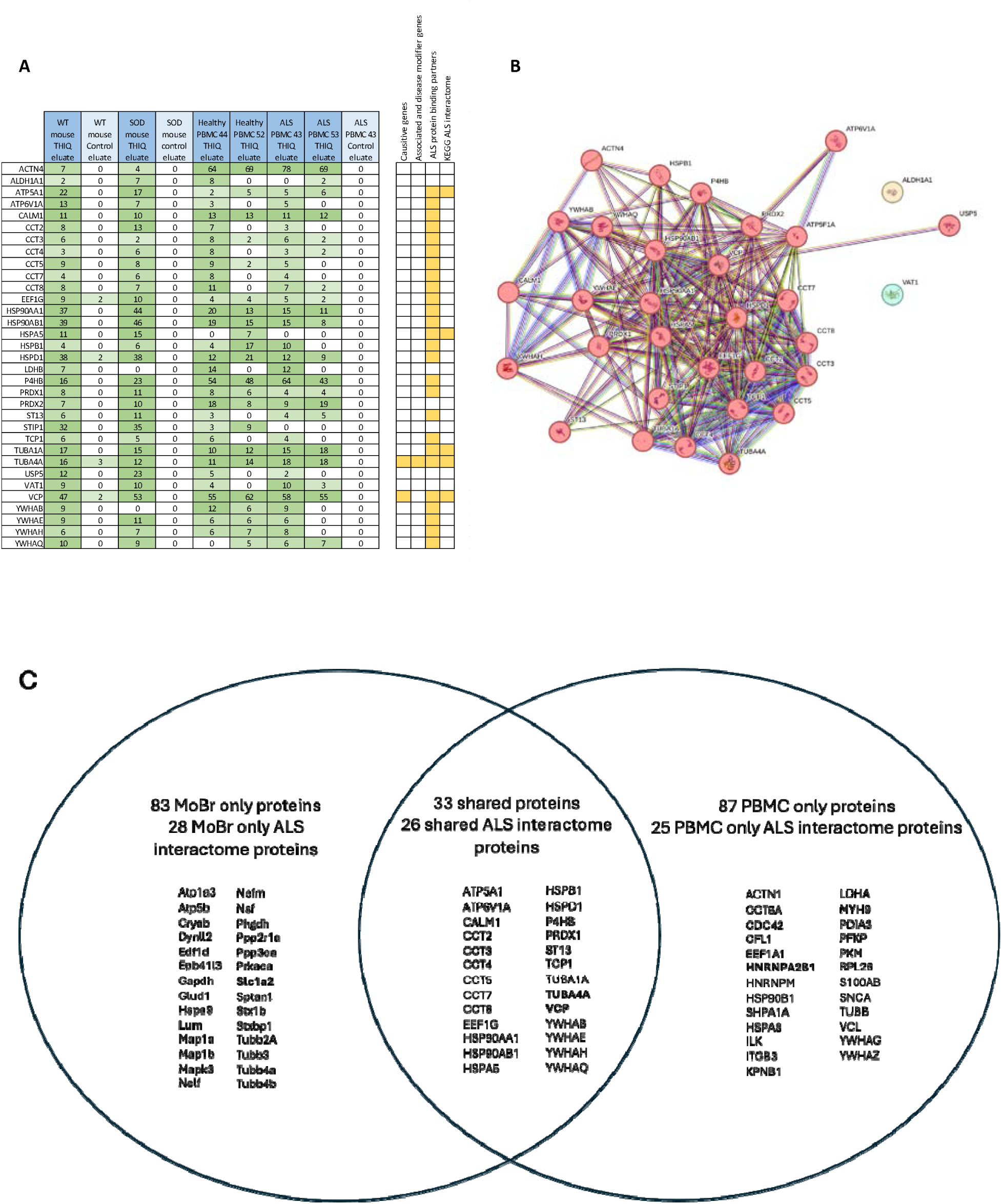
A. Spectral counts for proteins identified in THIQ drug resin eluate from MoBr and human PBMC extracts. Colors represent the number of spectral counts for a given protein in each sample, ranging from light green (low counts) to dark green (high counts). The right panel indicates “hits” for proteins present in the ALS interactome, where yellow is a hit. B. STRING database protein-protein interaction network with MCL clustering. Each circle (“node”) represents a protein, and each line (“edge”) represents a predicted interaction between two nodes. Different colors represent distinct clusters identified by the MCL algorithm, with red indicating the largest cluster and yellow and green representing clusters containing only a single protein. C. ALS interactome proteins identified by MS-MS are shown in venn diagram. Causative and disease modifier proteins are in **bold**.

MCL clustering analysis was performed on the 33 shared proteins using the STRING database, which integrates information from various databases cataloging known and predicted protein-protein interactions. The analysis generated groups called “clusters,” characterized by a higher density of predicted associations within the clusters compared to those between clusters. Notably, 31 out of the 33 proteins formed a single, highly interconnected cluster, suggesting potential functional relationships between them. Two proteins, VAT1 and ALDH1A1, were excluded from this cluster, as there are no predicted interactions between these proteins and the other 31 proteins (**Figure 2B**). These findings are summarized in **Figure 2C**.

From the MS-MS analysis it was clear that the multi-protein complex isolated from PBMCs on the ALS-active drug resin appears related to that observed in MoBr, based on the 27 shared proteins identified that are in the ALS protein interactome observed from both sources. However no clear differences between healthy and ALS patients were observed by MS-MS.

Recognizing that MS-MS analysis as carried out here can fail to identify specific proteins for a number of reasons, including covalent post-translational modifications (PTMs) that alter peptide mass, or result in failure of peptide cleavage with trypsin and/or ionization, we analyzed the DRAC bound material by WB to identify particular proteins of interest. WB has its own limitations, partially but not completely overlapping with those of MS-MS. A shared limitation with MS-MS is that a covalent post-translational can mask an epitope making it undetectable by an epitope-specific antibody. Nevertheless, if a difference between healthy and ALS patients in a protein within the drug resin-bound target complex were identified either by WB or MS-MS, it could serve as a signature of the disease. From WB of the DRAC bound material two proteins were strikingly different between most healthy individuals studied and most ALS patients. First, p62/SQSTM1, a known regulator of autophagy, an important host defensive mechanism, was greatly diminished in ALS patient PBMC DRAC free drug eluates compared to that from most healthy individuals. Conversely, a 17kDa post-translationally modified fragment of RanGTPase was detected in ALS patients, but not in most healthy individuals, in DRAC free drug eluates. A representative example is shown in **Figure 3**.

**Figure 3.**
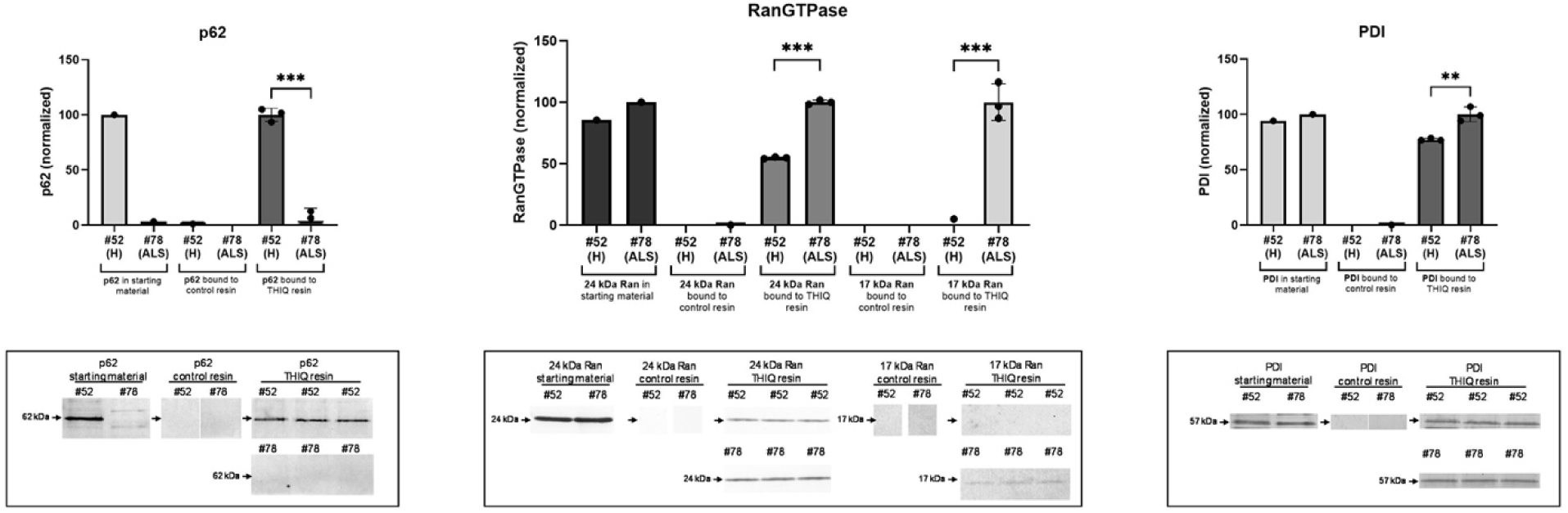
PBMC extracts were prepared from ALS patients (ALS) or healthy individuals (H) as described in methods and applied to drug resin versus control resin, washed and bound proteins analyzed by SDS-PAGE transferred to 0.2u PVDF membrane and WB for the indicated proteins (below) with quantitation of bands using Image J software (above). Shown is DRAC bound material from a healthy control (#52) and a sporadic ALS patient (#78) analyzed by WB for p62/SQSTM1, RanGTPase, and PDI. The highest value in each healthy/ALS graph pair is normalized to 100% for proportional comparison to the lower value.

Out of 13 ALS patients and 6 healthy controls assessed in a blinded fashion, all 13 ALS patients had a strikingly low values for p62 in the drug resin bound material. Of the 6 healthy controls, 2 were low. For Ran 17 kDa fragment, all 13 ALS patients had a higher value than the parallel reference healthy control. Among the 6 healthy controls, the two with low p62 also had high Ran 17 kDa fragment in the eDRAC bound material. In contrast to striking loss of p62 and appearance of RanGTPase 17kDa fragment, only a modest difference in the direct drug-binding protein, PDI, was observed between ALS patients and healthy controls.

Thus, taken together, the eDRAC signatures observed appear to have a sensitivity approaching 100% and a specificity of approximately 66% for ALS, albeit from a modest sample size. Further studies are needed to understand how these differences come about and what their significance is to the underlying pathophysiology of ALS. Nevertheless, together with the clinical picture, they suggest a basis for early detection of ALS using as an affinity ligand, a small molecule that is therapeutic for the disease in a diversity of animal models studied^14^. **Table 1** summarizes these findings together with key demographic features of the individuals studied. The eDRAC signature in PBMCs, like that observed previously in other cells and tissues, represent very small subsets of the total of the proteins in the target multi-protein complex (**Supplemental Figure 3**).

**Table 1.** 13 ALS patients and 6 healthy controls were analyzed over 4 separate experiments and the data tabulated, together with key demographic and clinical history information. For each experiment the highest value p62 was set at 100% and the others scaled proportionally. Likewise, in each experiment the highest value 17kDa Ran fragment was set at 100% and the others scaled proportionally. The quantified WB band value is also shown in parentheses. Healthy individuals are noted in green, and ALS patients in red, except for the two healthy individuals displaying the ALS phenotype, whose values are in black.

|  | % p62 | % 17kDa Ran Fragment | sex | age at onset | age dx | age at presentation | date of death | disease course | sample date (mo/day/yr) |
| --- | --- | --- | --- | --- | --- | --- | --- | --- | --- |
| #52 (healthy) | 100 (650) | 1 (2) | F |  |  | 41 |  |  | 7/31/18 |
| #40 (ALS) | 71 (462) | 5 (12) | M | 68 | 71 | 72 | N/A | LMN predominant - early bulbar/trach slow, progressive course | 1/24/19 |
| #50 (ALS) | 30 (193) | 12 (32) | M | 46 | 47 | 59 | 10/21/22 | Generalized slowly progressive disease | 7/31/18<br>1/24/19 |
| #51 (ALS) | 64 (419) | 8 (22) | M | 38 | 39 | 40 | 3/1/23 | UMN predominant generalized disease with bulbar compromise | 7/31/18 |
| #68 (ALS) | 3 (16) | 100 (261) | F | 78 | 79 | 79 | N/A | primary bulbar presentation with mostly UMN features | 2/19/19 |
| #54 (ALS) | 6 (41) | 76 (197) | M | 51 | 52 | 52 | 2/5/19 | rapidly progressive, bulbar involving disease | 8/7/18 |
| #55 (healthy) | 10 (63) | 86 (224) | F |  |  | 55 |  |  |  |
| #52 (healthy) | 80 (645) | 0 (0) | F |  |  | 41 |  |  | 7/31/18 |
| #45 (ALS) | 13 (64) | 61 (63) | M | 42 | 43 | 43 | N/A |  | 5/22/18<br>2/28/19 |
| #78 (ALS) | 6 (35) | 100 (103) | F | 45 | 46 | 46 | 10/7/20 | LMN predominant, generalized disease | 5/28/19<br>7/18/19 |
| #52 (healthy) | 99 (700) | 8 (8) | F |  |  | 41 |  |  | 7/31/18 |
| #87 (healthy) | 100 (706) | 5 (5) | M |  |  | 36 |  |  | 12/19/19 |
| #52 (healthy) | 100 (710) | 0 (0) | F |  |  | 41 |  |  | 7/31/18 |
| #43 (ALS) | 12 (85) | 100 (44) | F | 68 | 71 | 72 | 12/19/18 | UMN predominant; generalized disease with significant bulbar compromise | 12/19/18 |
| #44 (healthy) | 15 (105) | 27 (12) | F |  |  | 74 |  |  |  |
| #46 (ALS) | 54 (385) | 11 (5) | M | 53 | 54 | 54 | N/A | bulbar onset then generalized disease | 5/23/18 |
| #47 (healthy) | 56 (400) | 7 (3) | M |  |  | 58 |  |  |  |
| #53 (ALS) | 0 (0) | 68 (30) | F | 67 | 68 | 71 | 2/12/19 | limb onset generalized disease, possible concurrent Parkinson's disease | 7/31/18 |
| #26 (healthy) | 100 (558) | 0 (0) | M |  |  | 64 |  |  | 1/22/18 |
| #34 (ALS) | 1 (3) | 7 (2) | M | 40 | 42 | 42 | N/A | monomelic presentation VERY slow LMN generalization, bulbar sparing | 2/15/18 |
| #69 (ALS) | 2 (13) | 100 (24) | M | 80 | 82 | 82 | 5/8/21 | LMN predominant, generalized disease | 2/19/19 |
| #70 (ALS) | 13 (71) | 7 (2) | M | 42 | 42 | 43 | N/A | generalized progressive disease | 2/28/19 |

Next, photocrosslinking was carried out with PBMC extracts to which was added a drug analog in which the attachment to resin was replaced by attachment to a biotin and diazirine moiety such that exposure to UV light results in a covalent linkage to the drug binding protein(s). Under native conditions associated proteins of the entire multi-protein complex can be precipitated with streptavidin beads. If the sample is first denatured, then only the direct drug-binding protein will be precipitated with streptavidin beads under the conditions described. Figure 4 demonstrates that PDI is the direct drug-binding protein. It should be noted that P4HB found by MS-MS is the gene name for the same PDI family member detected by the PDI antibody by WB.

**Figure 4.**
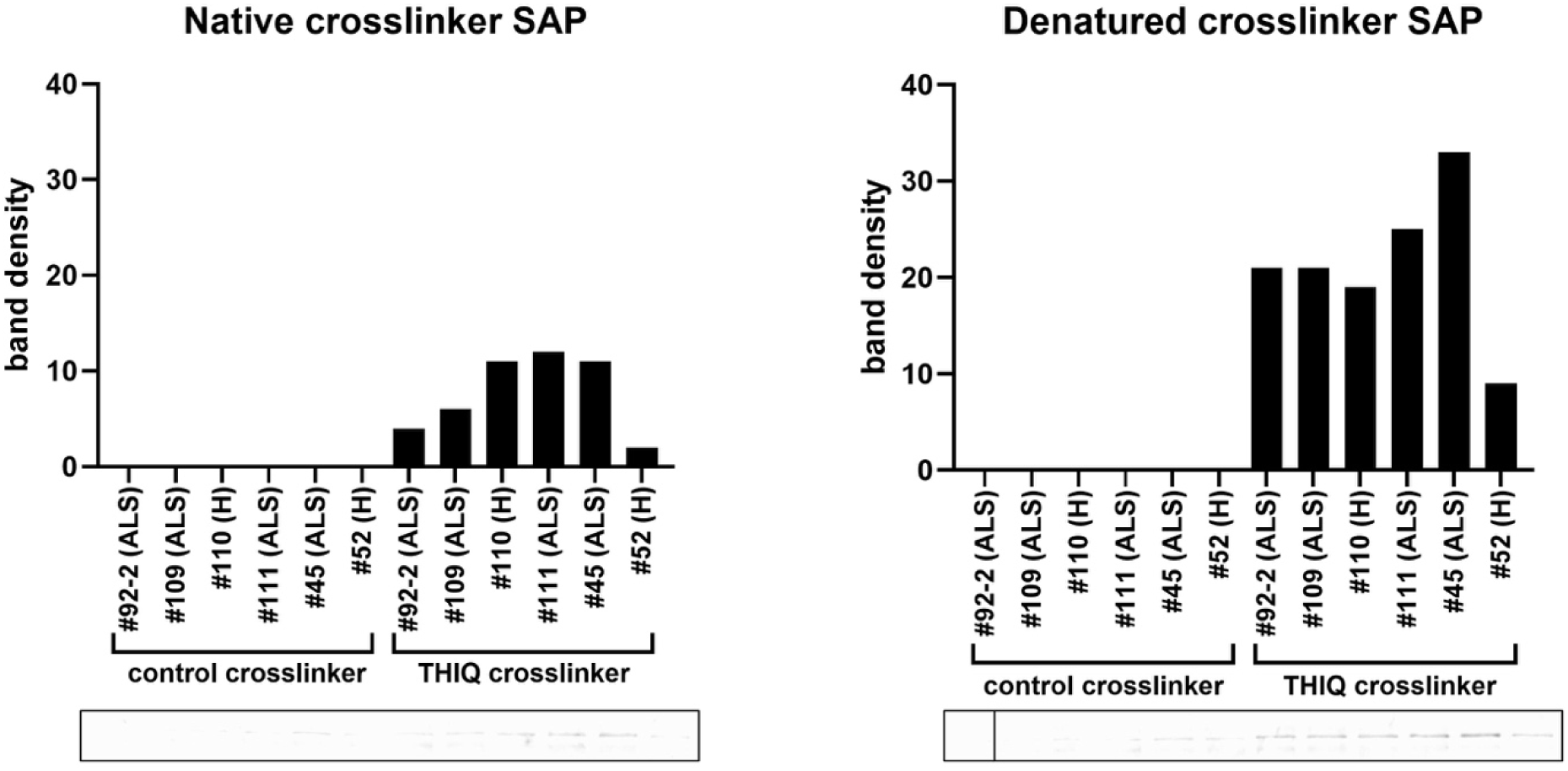
Quantitation of PDI in streptavidin precipitates from PBMC extracts prepared from ALS patients (ALS) or healthy individuals (H), under native vs denatured conditions after photocrosslinking with PAV-073 analog (THIQ) or control crosslinker lacking the drug ligand. Both control and THIQ crosslinkers contain biotin and diazirine as described in methods and previously^31^.

We then assessed whether the ALS patient vs healthy control correlations involving p62 and Ran 17kDa fragment, as shown in **Figure 3** and **Table 1**, were accentuated over time with progression of ALS. As can be seen in **Figure 5**, with progression of disease, the amount of PBMC p62 bound to the resin, which is much higher in healthy individuals (panel A), diminishes with advancement of ALS. In parallel, the amount of the RanGTPase 17kDa fragment, which is missing in PBMCs from healthy individuals, increases with ALS disease progression.

**Figure 5.**
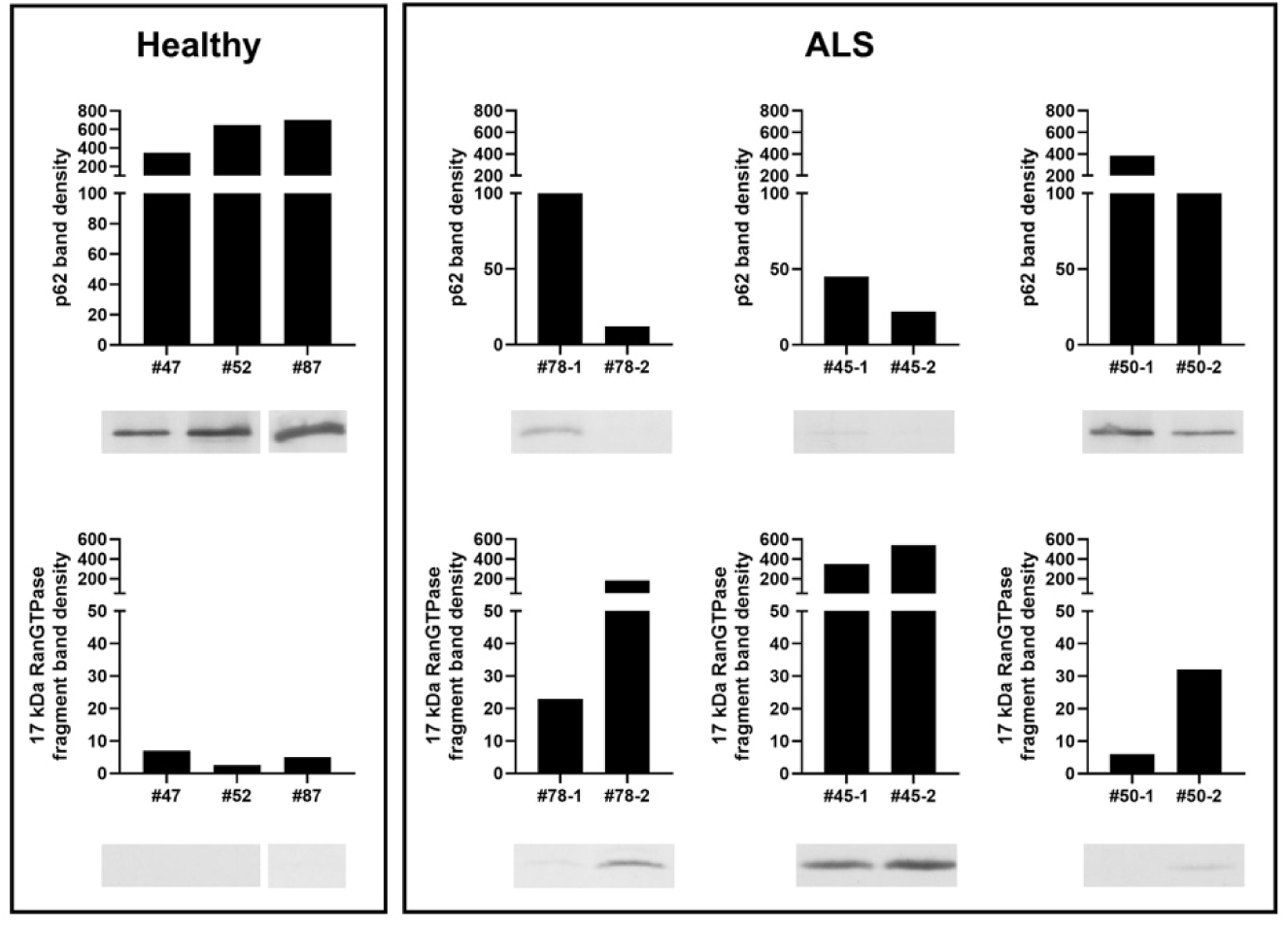
PBMC extracts were prepared as previously, applied to drug resins and bound proteins analyzed by WB for specific proteins of interest. Bands corresponding to the correct molecular weights were quantified. Blood samples from three ALS patients were assessed over time for both p62 and RanGTPase 17kDa fragment bound to DRAC. Timing of blood draws were 6 weeks apart for patient 78, 6 months apart for patient 50, and 9 months apart for patient 45.

Notably, the diagnosis of ALS was made in pt 78 very early in her disease progression. She was able to drive herself to the clinic and walk in with a cane, at the time that the initial blood sample was taken, which nevertheless revealed the dual signatures of falling p62 and appearance of Ran 17kDa fragment, in contrast to parallel analysis of a healthy control. As will be seen these features appear to be exacerbated with disease progression.

If, as we suggest, ALS represents a breakdown in homeostasis where protective mechanisms to reboot a cellular operating system are ineffective, certain predictions follow. For one, there may be a backup mechanism activated in an attempt to correct the failure of the first line of defense, although by definition, in patients who progress to ALS, that backup has failed to correct the problem. By analogy, when your computer is frozen (with appearance of the “colored wheel of doom”), simply attempting to undo the key stroke that caused it to freeze will not be effective. Instead, it is necessary to reboot the operating system. Perhaps assembly modulators provide precisely such a biochemical reboot of the cellular operating system. In view of the distinctive ALS-associated appearance of a 17kDa fragment of RanGTPase, as noted in **Figure 5** above, degradation of RanGTPase, seemed like a plausible backup feedback response. This backup feedback response would be engaged if less draconian feedback methods failed.

A second prediction is that if, as we hypothesize, assembly modulation is a fundamental mechanism of regulation of gene expression, and its activation by these compounds is therapeutic for ALS, treatment of whole blood from ALS patients with the drugs should result in cessation of the backup RanGTPase degradation feedback loop, upon “reboot” of homeostatic control.

To test these hypotheses, we took blinded blood samples from ALS patients and healthy controls and treated the whole blood with two different, structurally unrelated ALS therapeutic assembly modulator compounds versus vehicle control (DMSO). Since RanGTPase is implicated in ALS pathophysiology and in the PBMC signature presented in Figures 3 and 5 we chose to assess a readily measurable potential feedback response: degradation of total RanGTPase, even though only a small subfraction of RanGTPase is hypothesized to be involved in ALS pathophysiology. We assessed the stability of RanGTPase with and without 72 hrs of assembly modulator treatment. After the 72 hr incubation of whole blood, RanGTPase stability in extracts was assessed in a 1hr incubation (**Figure 6**). In some samples, RanGTPase was observed to be stable over the course of a 1 hour incubation. These proved to be the healthy controls, when unblinded. However in blood from all of the ALS patients, the levels of RanGTPase after the 1hr incubation of whole blood extracts were greatly diminished, consistent with what would be expected for degradation of all RanGTPase as an extreme feedback response to the progression of ALS. Upon treatment with either of advanced analogs of two ALS assembly modulator chemotypes (called 073 and 667) for 72hrs, a striking diminution of RanGTPase degradation on subsequent 1hr extract incubation was observed, with no change in the level of RanGTPase observed in healthy subjects.

**Figure 6.**
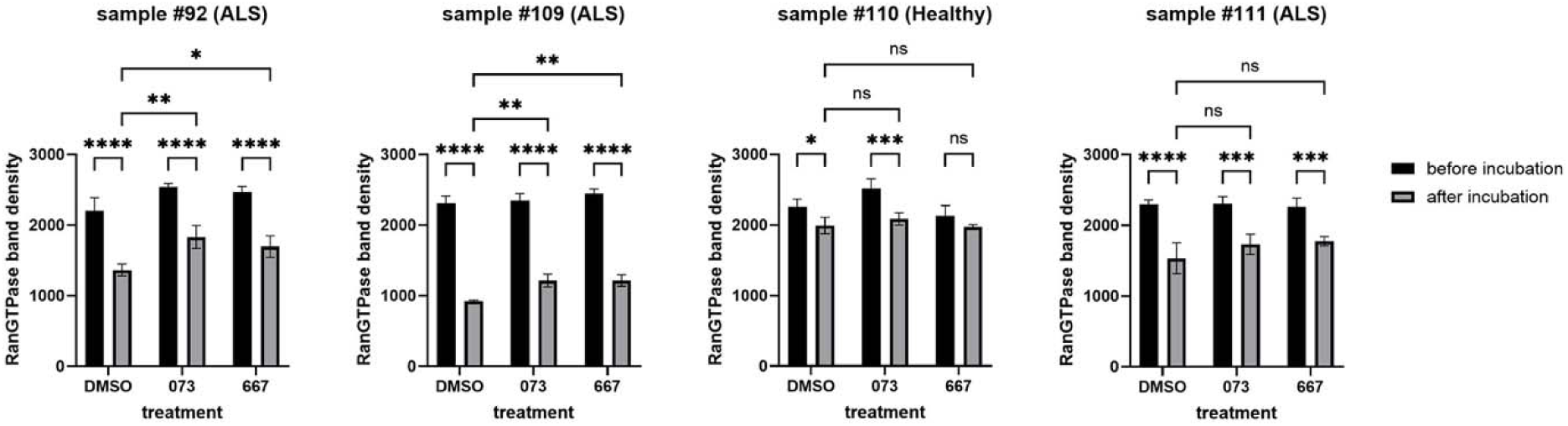
10mls of whole blood from ALS patients or healthy controls was collected under an IRB-approved protocol and shipped overnight at room temperature in EDTA-anticoagulated (purple top) tubes without any patient-specific confidential information. An aliquot of whole blood was taken, incubated in 1% SDS for one hour at 37°C, prepared for SDS-PAGE, and set aside. Another aliquot of whole blood was mixed 1:1 with glucose-supplemented MEM and incubated for 72hrs at 37°C in the presence of vehicle or protein assembly modulator drugs 073 and 667, at the end of which samples were processed as previously described, and analyzed by SDS-PAGE. Black bars refer to the sample prepared for SDS-PAGE immediately, while gray bars refer to the samples incubated prior to preparation for SDS-PAGE.

## Discussion

The studies presented here demonstrate the presence, by energy-dependent DRAC and photocrosslinking, of a multi-protein complex in PBMCs that is related to that observed previously by the same methods in Tg ALS SOD G93A model MoBr and ALS patient-derived fibroblasts (PDFs) that display both TDP-43 mislocalization and stress-induced agregation^14^. As observed previously for the drug target in MoBr and PDFs, the multi-protein complex drug target in PBMCs has a small subset of PDI as the direct drug-binding protein. The complex present in ALS patient PBMCs also shares with the target in MoBr the presence of 26 proteins implicated in the ALS interactome (Figure 2C). The precise role of this multi-protein complex drug target in ALS pathophysiology remains to be determined. However, based on the efficacy of structurally unrelated ALS assembly modulators in various cellular and animal models, we hypothesize that the drug target serves a role in maintenance of homeostasis, whose loss results in ALS. The presence of p62 in the complex in PBMCs from healthy individuals, and its striking diminution in ALS patient PBMCs, exacerbated with disease progression (Figure 5), are consistent with this hypothesis.

Likewise, the appearance of a 17kDa post-translationally modified form of RanGTPase in the drug target isolated from ALS patient PBMCs (Figure 3 and Table 1) suggests a complex biology connected to TDP-43 mislocalization, in view of the crucial role of RanGTPase in nucleo-cytoplasmic transport. These changes appear very early in ALS, as suggested by their detection in patient #78 prior to onset of significant disability (Figure 4). In the three cases where a second sample was assessed 6 weeks to 9 months after the first blood draw, both signatures were seen to be exacerbated with disease progression. These signatures had a sensitivity of 100%, detected in PBMCs of all 13 sporadic ALS patients studied, and a specificity of 66% for distinguishing ALS patients from healthy individuals. While a larger number of cases will need to be assessed, this early degree of discrimination, using as an affinity ligand a drug shown therapeutically active in a diversity of animal models, is promising as a novel path to an ALS biomarker and highly relevant to ALS therapeutics.

This target multi-protein complex is implicated in ALS for four reasons. First, the drug that served as the ligand to bind and isolate the target is therapeutic in a wide range of ALS cellular and animal models^31^. Second, the composition of the target multi-protein complex includes a number of proteins of the ALS interactome, 26 of which are shared with the corresponding complex in MoBr. Third, the isolated target from ALS patient PBMCs is altered in protein composition compared to that from most healthy controls by two metrics: diminution of p62/SQSTM1 and appearance of 17kDa fragment of Ran GTPase. Finally, separate from the specific target in PBMCs, a profound degradation of total RanGTPase is observed in whole blood from ALS patients but not in healthy controls, and is substantially rescued after 72hrs of treatment with ALS-active assembly modulators.

We interpret these findings to suggest that ALS is, fundamentally, a disease of disrupted homeostasis, for which assembly modulators may be therapeutic. By analogy to another therapeutic area where assembly modulation treatment may be effective^16^, it is notable that the vast majority of individuals don’t have cancer for most of their lives. However, a small number will develop cancer, at least in part due to failure of homeostatic host defenses. So also host defenses protect most of us most of the time from the consequences of various central nervous system (CNS) insults, including those causing ALS. Host defenses, including restoration of homeostasis, usually result in resolution of a CNS insult. Sometimes however, those host defenses fail. As a result, homeostasis is not restored, and progression to motor neuron death is accelerated. It seems reasonable to hypothesize that when initial feedback loops fail to restore homeostasis, more and more severe feedback responses are triggered. Thus, mass degradation of RanGTPase, as observed here in whole blood from ALS patients (Figure 6), may represent a final desperate attempt to restore defective nucleocytoplasmic transport that results in TDP-43 mislocalization. Autophagy, in part mediated by p62/SQSTM1, is an example of a biochemical reboot of the cellular operating system. When the reboot fails, e.g. in part due to loss of p62 from the dysfunctional target, the consequence in a subset of cases may be ALS. This persistent deviation from homeostasis is a stress to which motor neurons appear highly sensitive, and eventually, succumb. A drug capable of restoring homeostasis is likely most effective before motor neurons have died, although perhaps synaptic plasticity could synergize allowing some degree of recovery, once homeostasis has been restored. Regardless, it is surely a truism of medicine that the earlier a disease is detected, the more effective most therapeutic interventions are likely to be. Hence the importance of early detection to enable early treatment, which the PBMC signatures noted here may provide.

The results of the whole blood experiments (Figure 6) are consistent with the hypothesis that the degradation of total RanGTPase observed, is an extreme feedback response to the inability more proximal and focused feedback loops to restore the target complex, comprised of a small subset of RanGTPase, back to the healthy state. Treatment with either of two structurally unrelated assembly modulators results in correction of the RanGTPase degradation observed in whole blood, to a significant extent. We hypothesize that treatment for a longer period would have completely restored RanGTPase levels to normal. Whether assembly modulator drug treatment serves to fully reboot the system, e.g. restore p62 to the target complex as has been observed for assembly modulation using a structurally unrelated molecule in a different therapeutic area^15^, remains to be determined and may vary on a case-by-case basis. Likewise, further studies are needed to determine whether periodic or ongoing drug treatment are required. Regardless, drug-induced changes in target composition, as implied by stabilization of RanGTPase in ALS patient blood treated with assembly modulators, is consistent with restoration of homeostasis. Whether this is reflective of concomitant changes in motor neurons that prevent further motor neuron loss, remains to be investigated.

Our findings are significant for a number of reasons. First, these studies provide a window into novel regulatory feedback loops involved in cellular homeostasis. The precise relationship of the target multi-protein complex in healthy individuals vs ALS patients remains to be more fully understood. One hypothesis is that the target multi-protein complex in healthy individuals is the “normal” assembly machine involved in integration of diverse events of homeostasis. These may include TDP-43 localization, in which RanGTPase likely plays a critical role, perhaps along with C9orf72 and other ALS-implicated gene products. Other relevant pathways, including autophagic removal of dysfunctional multi-protein complexes, are naturally integrated into the physiological feedback that collectively bring about homeostasis. ALS may be an example where a “stuck” pathway triggers more and more desperate feedback attempts to reboot the system without success, culminating in the mass degradation of RanGTPase observed in the whole blood experiment shown in Figure 6. In this model, the assembly modulator compound action can be thought of as serving as an external rather than internal reboot of the cellular operating system. A future prediction to be tested, is that longer-term treatment of blood results in complete normalization of RanGTPase – and restoration of p62/SQSTM1 -- in the target complex present in PBMCs. By virtue of having therapeutic small molecules as affinity ligands for target identification, the pathways discovered can be productively illuminated by future studies, including by fractionation and reconstitution of functional activities in extracts.

Second, these data, together with the studies shown previously, are consistent with the hypothesis that, despite the heterogeneity of ALS clinical presentation and time course of progression, there is a common underlying pathway that is the basis of the disease – and that assembly modulators are therapeutic at that underlying shared level. While we hypothesize that restoration of autophagy is a crucial part of this pathway, re-establishment of homeostasis likely involves far more than just restoration of autophagy. In this regards the involvement of miniscule subsets of the specific gene products found together in the target multi-protein complex and the role of protein degradation, is notable. These targets could not have been detected by powerful molecular biological tools such as CRISPR or siRNA knockdown studies, because of the heterogeneous roles of small subsets of many, if not most, of the proteins comprising the target multi-protein complex. Thus the concept of protein “moonlighting”^34,35^ may be more pervasive than is currently generally recognized. Conversely, the level of organization of gene expression accessed by protein assembly modulation may be therapeutic for diverse diseases in ways not generally appreciated today.

Finally, it should be noted that many if not most patients come to medical attention long after the most effective period of treatment is lost. Drugs to stop the disease process may still be effective, but rely on maintenance of the underlying framework of feedback loops to restore homeostasis, once disease progression is arrested. Sometimes however, those feedback loops fail and homeostasis is not restored, whether or not the primary disease process has been blocked. We hypothesize ALS is one such example, making it extremely difficult to treat. The methods described here provide a means of detecting ALS early, prior to onset of severe disability, when it is most treatable – and provide a potentially effective treatment in the form of the small molecule ligands themselves, that were used for disease detection. It is notable that pt 78 came to medical attention early in her clinical course, prior to serious disability, and yet, the signature changes (lowered p62 and elevated RanGTPase 17kDa fragment) were nevertheless observed, and sadly, progressed with worsening of ALS, culminating in her death, approximately 2 years after initial detection of the Ran 17kDa fragment as shown in Figure 5, together with the reciprocal change in p62/SQSTM1, seen exacerbated over time. To date, these findings have been observed in 13 of 13 ALS patients and are lacking in 4 of 6 healthy controls, by the methods described. While the specificity of these signatures are relatively low, seen in 2 of 6 healthy controls, the high sensitivity implied by detection in 13 of 13 ALS patients, together with the clinical history, make this diagnostic approach promising. PBMCs are a readily accessible tissue. Used as affinity ligands, these small molecules could enable early diagnosis of ALS, prior to extensive motor neuron death, when treatment would be more effective. Remarkably, the same compounds, or their close analogs, may prove to be effective ALS therapeutics.

## Methods

### Preparation of PBMC extracts^36^

Whole blood collected in purple-top EDTA tubes from healthy individuals or ALS patients under Loma Linda Medical Center IRB# 5160136 was shipped at room temperature by overnight courier to the San Francisco lab where it was diluted 1:1 with phosphate-buffered saline and layered on ficoll and centrifuged for 15 minutes at 1,000 xg at room temperature with brake off. PBMCs were collected at the interface, diluted with PBS 1:1 and centrifuged at 600 xg /15 minutes. The supernatant was aspirated and the pellet dissolved in p-body buffer (PBB) consisting of 10mM Tris pH 7.5, 10mM NaCl, 6mM MgAc, 1mM EDTA, and 0.35% Triton-X-100 and centrifuged at 4oC 10,000xg/10min. The 10kxg supernate was collected, aliquoted and flash frozen in liquid nitrogen and stored at −80°C until use.

### Assembly modulator compounds

Each assembly modulator chemotype is structurally quite different from one another, depending on the target to which it is directed, the allosteric site involved, etc. The two ALS-active assembly modulator chemotypes used here, typified by compounds PAV-073 and PAV-667 are compared for drug-like properties in Table 2. The structure of PAV-073, a member of the THIQ chemotype, and its modification to allow its use as an affinity ligand for drug resin affinity chromatography or photocrosslinking, is described in reference 14.

**Table 2.** Two structurally unrelated (Tanimoto similarity coefficient <20 %) assembly modulator chemotypes used in the experiments shown in Figure 6 are compared for drug-like properties.

| Analog representative of chemotype | Molecular weight (g/mol) | log P | H-bond donors | H-bond acceptors | Topological polar surface area (Å²) | Tanimoto similarity coefficient |
| --- | --- | --- | --- | --- | --- | --- |
| PAV-073 | 443.543 | 4.8 | 1 | 5 | 49 | <20% |
| PAV-667 | 441.915 | 6.4 | 0 | 7 | 73.7 | <20% |

### Drug Resin affinity chromatography

A variation on the theme of methods described in previous publications was used here^14–16^. For brain from wildtype or SOD1 mutant animals, tissue was homogenized in cold low salt buffer (10mM HEPES pH 7.6, 10mM NaCl, 1mM MgAc with 0.35% Tritonx100) then centrifuged at 10,000 xg for 10 minutes at 4°C. The post-mitochondrial supernatant was adjusted to a concentration of approximately 10 mg/ml and equilibrated in a physiologic column buffer (50 mM HEPES ph 7.6, 100 mM KAc, 6 mM MgAc, 1 mM EDTA, 4mM TGA). For DRAC of both MoBr and human PBMC extracts, the 10k sup samples being compared were first normalized either by absorbance at 280nm or by Coomassie brilliant blue or silver staining pattern. Samples were then supplemented with an energy cocktail (to a final concentration of 1mM rATP, 1mM rGTP, 1mM rCTP, 1mM rUTP, and 5 µg/mL creatine kinase) in the presence of physiological salts (50mM HEPES pH 7.6, 100mM Kac, MgAc 5mM, 1mM DTT) and incubated for one hour at either 4° C or 22° C degrees on 30 µl of affigel resin coupled to THIQ compound or a 4% agarose matrix (control). The input material was collected after flow through the column and the resin was then washed with 3 ml column buffer. The resins were eluted for 2 hours then overnight at 22°C then 4°C in 100µl column buffer containing 100µM of the cognate compound. Eluates were run on WB or sent for MS-MS for analysis.

### Chemical photocrosslinking

A variation of protocols as published previously^14^ was used. Extract from MoBr were prepared as described above then adjusted to a protein concentration of approximately 3 mg/ml in column buffer (Hepes pH 7.5 50mM, Kac 100mM, MgAc 5mM, DTT or TGA 1mM) containing 0.1% triton. 1% DMSO or 100µM PAV-073 was added to 6µl of extract, then 3µM of PAV-073 photocrosslinker or a negative control crosslinker (comprising of the biotin and diazirine moieties without compound) were added. The extract was incubated for 20 minutes then exposed to UV at 365nM wavelength for 10 minutes then left on ice for one hour. After crosslinking, samples were divided in two 20 µl aliquots and one set was denatured by adding 20 µL of column buffer 4µl of 10% SDS, 0.5 µl 1M dithiothreitol (DTT), and heating to 100°C for 5 minutes. Both native and denatured aliquots were then diluted in 800 µl column buffer containing 0.1% triton. 5 µl of magnetic streptavidin beads (Pierce) were added to all samples and mixed for one hour at room temperature to capture all biotinylated proteins and co-associated proteins. Samples were placed on a magnetic rack to hold the beads in placed and washed three times with 800 µl of column buffer containing 0.1% triton. After washing, beads were resuspended in 80 µl of gel loading buffer containing SDS and analyzed by SDS-PAGE followed by transfer to PVDF membrane for WB.

### WB Procedure

SDS-PAGE gels were transferred in Towbin buffer (25mM Tris, 192mM glycine, 20% w/v methanol) to polyvinylidene fluoride membrane, blocked in 1% bovine serum albumin (BSA) in PBS, incubated overnight at 4°C in a 1:1 dilution of 100µg/mL affinity-purified primary rabbit IGG to PDI (Abcam 96744) in 1% BSA in PBS containing 0.1% Tween-20 (PBST). Membranes were then washed twice in PBST and incubated for two hours at room temperature in a 1:5,000 dilution of secondary anti-rabbit or anti-mouse antibody coupled to alkaline phosphatase in PBST. Membranes were washed two more times in PBST then incubated in a developer solution prepared from 100 µL of 7.5 mg/mL 5-bromo-4-chloro-3-indolyl phosphate dissolved in 60% dimethyl formamide (DMF) in water and 100µl of 15 mg/ml nitro blue tetrazolium dissolved in 70% DMF in water, adjusted to 50mL with 0.1 Tris (pH 9.5) and 0.1 mM magnesium chloride. Membranes were scanned and the integrated density of protein band was measured on ImageJ. Averages and the standard deviation between repeated experiments were calculated and plotted on Microsoft Excel. Primary antibodies for RanGTPase (Abcam ab31118), p62/SQSTM1 (Abcam, ab56416), were used at 1:1,000 dilution by the same procedure. Secondary goat antibodies to mouse (Abcam ab97020) or rabbit (Abcam ab6722) were used at 1:5,000 dilution.

### Tandem mass spectrometry

Samples were processed by SDS PAGE using a 10% Bis-tris NuPAGE gel with the 2-(N-morpholino)ethanesulfonic acid buffer system. The mobility region, including all proteins, was excised and washed with 25 mM ammonium bicarbonate followed by 15mM acetonitrile. Samples were reduced with 10 mM dithoithreitol and 60° C followed by alkylation with 50 mM iodoacetamide at room temperature. Samples were then digested with trypsin (Promega) overnight (18 hours) at 37° C then quenched with formic acid and desalted using an Empore SD plate. Half of each digested sample was analyzed by LC-MS/MS with a Waters NanoAcquity HPLC system interfaced to a ThermoFisher Q Exactive. Peptides were loaded on a trapping column and eluted over a 75 µM analytical column at 350 nL/min packed with Luna C18 resin (Phenomenex). The mass spectrometer was operated in a data dependent mode, with the Oribtrap operating at 60,000 FWHM and 15,000 FWHM for MS and MS/MS respectively. The fifteen most abundant ions were selected for MS/MS.

Data was searched using a local copy of Mascot (Matrix Science) with the following parameters: Enzyme: Trypsin/P; Database: SwissProt Human (conducted forward and reverse plus common contaminants); Fixed modification: Carbamidomethyl (C) Variable modifications: Oxidation (M), Acetyl (N-term), Pyro-Glu (N-term Q), Deamidation (N/Q) Mass values: Monoisotopic; Peptide Mass Tolerance: 10 ppm; Fragment Mass Tolerance: 0.02 Da; Max Missed Cleavages: 2. The data was analyzed by spectral count methods.

#### Diagnosis of ALS

A diagnosis of ALS was made in all patients by a combination of their neurological exam and their electrodiagnostic testing. The examination was required to reveal signs of upper and lower motor neuron pathology. Although the phenotypic spectrum of ALS is quite variable, upper and lower motor neuron signs were represented in all patients and a brief phenotypic description is included in Table 1. Electrodiagnostic data included multifocal evidence of denervation and reduced motor neuron recruitment.

#### Whole blood incubation and analysis

100µl aliquots of EDTA-anticoagulated whole blood is dispensed into 24 well plates containing 100µl of room temperature 80% PBS with 10mM glucose and then treated with DMSO to 0.01% or drug in DMSO to the same dilution with final concentration of 5µM drug. After 72hrs samples are collected and adjusted to 1% SDS. One aliquot was prepared in gel loading buffer as starting material. Another aliquot added to energy cocktail (1mM rNTPs with 0.2µg/ml CK, 50mM Hepes pH 7.6 100mK KAc, 6mM MgAc and 4mM TGA) at total volume 100µl and incubated at 37°C for 1hr, and then prepared in gel loading buffer. Samples were analyzed by SDS-PAGE and WB for full length RanGTPase.

### Statistics

Graphs were generated and statistical analysis were preformed using GraphPad prism version 10.4.1. For Figure 3, unpaired t-tests were conducted to assess statistical significance in the amount of protein bound to the THIQ resin for each starting material. In figure 6, a 2way ANOVA was used, followed by Tukey’s multiple comparisons tests. P-values of selected comparisons were shown on the graph, including changes in RanGTPase levels before and after treatment within each condition and differences in RanGTPase levels between DMSO and drug treatments post-treatment. Statistical significance is indicated as follows: nsa (P > 0.05), * (P ≤ 0.05), ** (P ≤ 0.01), *** (P ≤ 0.001), **** (P ≤ 0.0001).

### Ethical Approval

Whole blood collected in purple-top EDTA tubes from healthy individuals or ALS patients under Loma Linda Medical Center IRB# 5160136

---

**Supplemental Figure 1.**
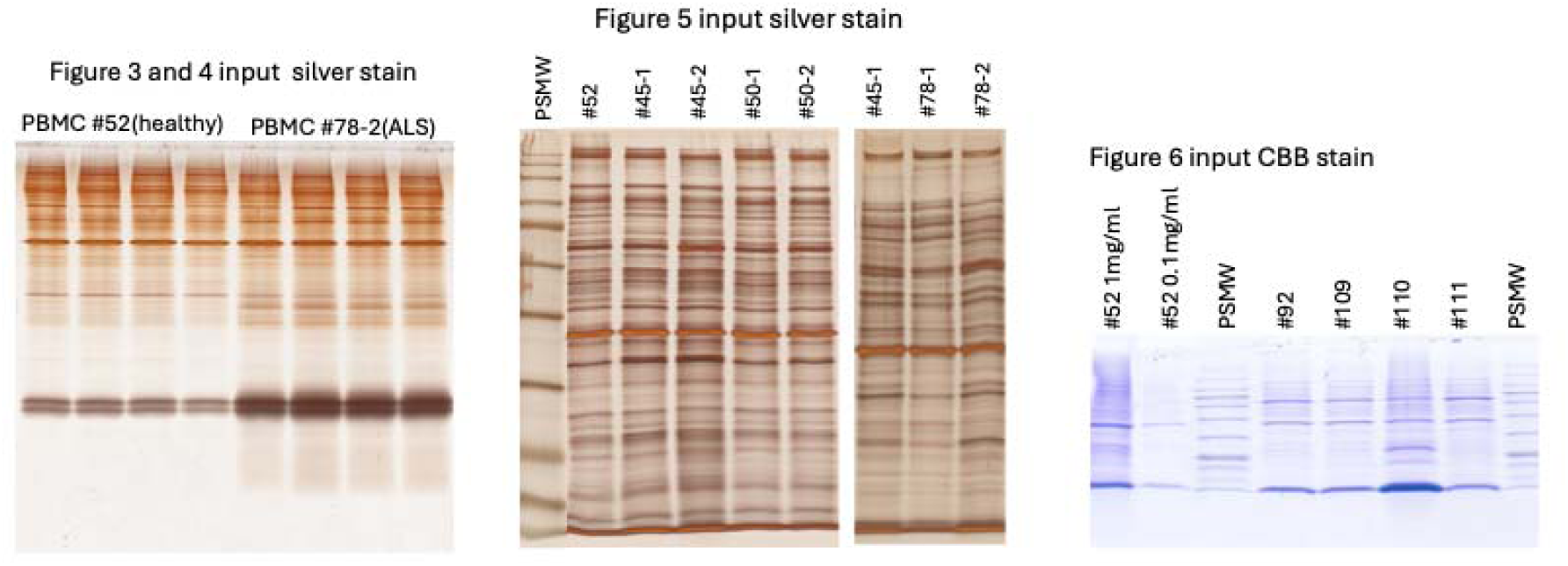
Input extracts for Figures 2-6. Silver stain (left and middle panels) and CBB stain (right panel) of input extracts used for DRAC (Figure 3 and 5), photocrosslinking (Figure 4), amd RanGTPase degradation (Figure 6) assays.

**Supplemental Figure 2.**
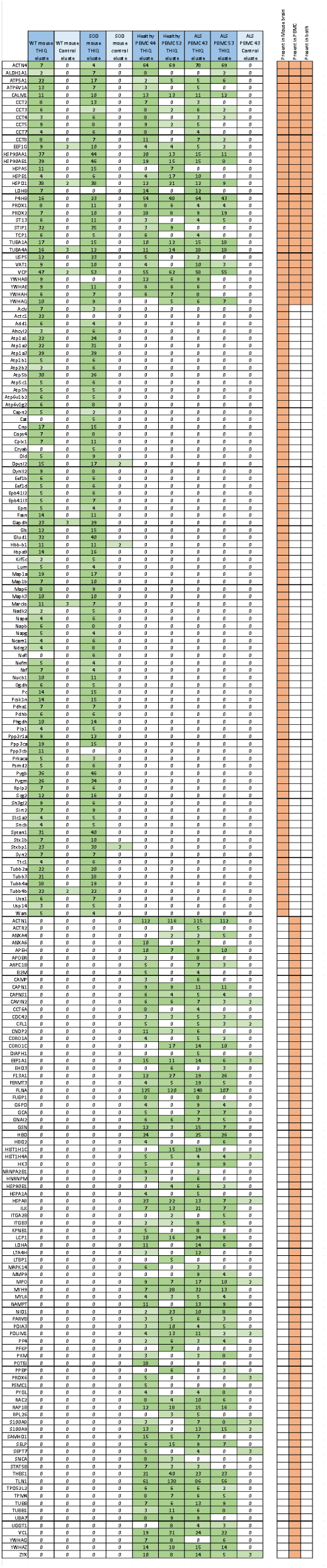
Spectral counts for entire dataset of MoBr and PBMC THIQ resin eluates. Colors represent the number of spectral counts for a given protein in each sample, ranging from light green (low counts) to dark green (high counts). The right panel indicates whether a given protein was in one of both of the datasets, according to threshold values of >4 spectral counts in the drug eluate and <4 spectral counts in the control column eluate.

**Supplemental Figure 3.**
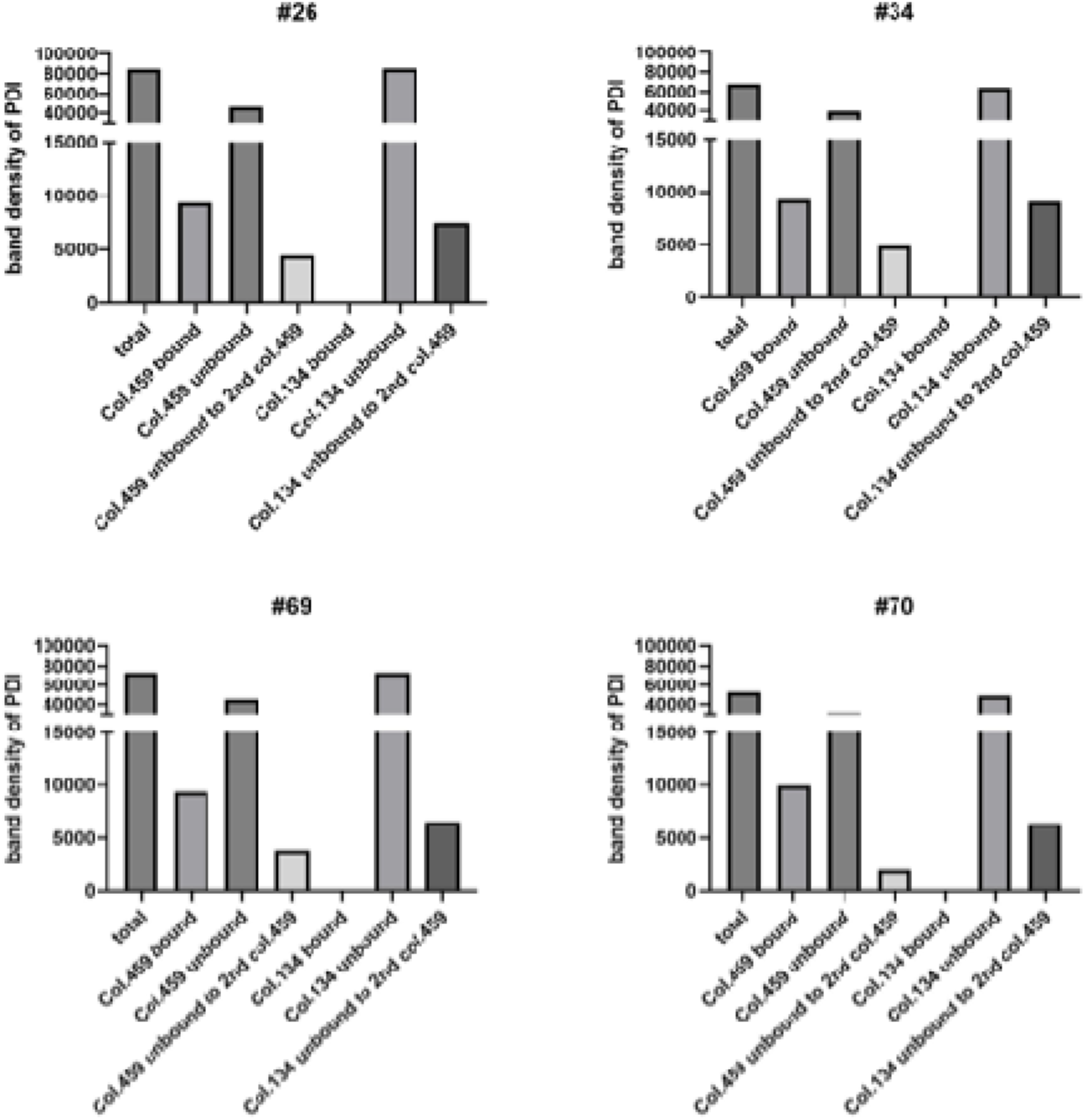
Confirmation that only a small subset of PDI is present in the DRAC-identified target and that very little additional pre-existing PDI is competent for formation of new target without *de novo* synthesis. PBMC extracts were prepared as previously, applied to drug resins and bound proteins analyzed by WB for PDI. The band corresponding to the correct molecular weight was quantified as shown. Samples from three ALS patients and one healthy control are shown. Column 459 represents the THIQ drug resin and column 134 the control drug resin. As can be seen, approximately 10% of PDI is in the complex bound to the drug resin in all cases, with no significant binding to the control resin in any case. The flowthrough from the first column (drug or control) was applied to a second copy of the drug resin, demonstrating substantial depletion from the first column and that, despite losses, a substantial fraction of the flow through from the control column remains competent to bind to the drug resin column.

## Abbreviations

ALS: Amyotrophic Lateral Sclerosis
BSA: bovine serum albumin
CFPSA: Cell-free protein synthesis and assembly
Col: column
Ctrl: control
CNS: Central Nervous System
DTT: dithiothreitol
DRAC: drug resin affinity chromatography
Dx: diagnosis
eDRAC: energy-dependent drug resin affinity chromatography
FTD: Fronto-Temporal Dementia
H: healthy individual
HEPES: 4-(2-hydroxyethyl)-1-piperazineethanesulfonic acid
IND: investigational new drug
LMN: lower moter neuron
MoBr: mouse brain
MS-MS: tandem mass spectrometry
ns: non-significant
PBMC: Peripheral blood mononuclear cell
PBS: phosphate-buffered saline
PDF: patient-derived fibroblast
PDI: protein disulfide isomerase
PTM: post-translationally modified
PSMW: Prestained molecular weight markers
PVDF: polyvinylidene difluoride
SAP: streptavidin precipitation
SDS-PAGE: polyacrylamide gel electrophoresis in sodium dodecyl sulfate
TDP-43: Transactive Response DNA-binding Protein 43
TGA: thioglycolic acid
TPP: target product profile
UMN: upper motor neuron
WB: western blot
*: p value < 0.05
**: p value < 0.01
***: p value < 0.001
****: p value < 0.0001

## Dataset

The relevant dataset has been submitted to the DRYAD database and can be accessed upon publication at the following: DOI: 10.5061/dryad.0rxwdbsbm

## Competing Interests

VRL is CEO and AFL COO of Prosetta Biosciences

## Funding Statement

This work was supported by Target ALS (grant# IL-2023-C4-L1), DOD CDMRP (grant #. W81XWH2210721), and Prosetta Biosciences.

## Acknowledgements

We thank David Hanzel and the anonymous reviewers for careful reading of the manuscript and numerous helpful suggestions.

## Notes

### Competing Interest Statement

VRL is CEO and AFL is COO of Prosetta Biosciences

### Summary of Updates

The manuscript has been revised to respond to reviewer comments and various typographical errors and omissions have been corrected.

## References

1. Al-Chalabi, A. & Hardiman, O. The epidemiology of ALS: a conspiracy of genes, environment and time. Nat. Rev. Neurol. 9, 617–628 (2013).

2. Chiò, A. et al. Prognostic factors in ALS: A critical review. Amyotroph. Lateral Scler. Off. Publ. World Fed. Neurol. Res. Group Mot. Neuron Dis. 10, 310–323 (2009).

3. Rizea, R. E., Corlatescu, A.-D., Costin, H. P., Dumitru, A. & Ciurea, A. V. Understanding Amyotrophic Lateral Sclerosis: Pathophysiology, Diagnosis, and Therapeutic Advances. Int. J. Mol. Sci. 25, 9966 (2024).

4. Hernan-Godoy, M. & Rouaux, C. From Environment to Gene Expression: Epigenetic Methylations and One-Carbon Metabolism in Amyotrophic Lateral Sclerosis. Cells 13, 967 (2024).

5. Cabrera-Rodríguez, R. et al. TDP-43 Controls HIV-1 Viral Production and Virus Infectiveness. Int. J. Mol. Sci. 24, 7658 (2023).

6. Gandhi, J. et al. Protein misfolding and aggregation in neurodegenerative diseases: a review of pathogeneses, novel detection strategies, and potential therapeutics. Rev. Neurosci. 30, 339–358 (2019).

7. Grad, L. I., Rouleau, G. A., Ravits, J. & Cashman, N. R. Clinical Spectrum of Amyotrophic Lateral Sclerosis (ALS). Cold Spring Harb. Perspect. Med. 7, (2017).

8. Chou, C.-C. et al. TDP-43 pathology disrupts nuclear pore complexes and nucleocytoplasmic transport in ALS/FTD. Nat. Neurosci. 21, 228–239 (2018).

9. Baradaran-Heravi, Y., Van Broeckhoven, C. & van der Zee, J. Stress granule mediated protein aggregation and underlying gene defects in the FTD-ALS spectrum. Neurobiol. Dis. 134, 104639 (2020).

10. Gasset-Rosa, F. et al. Cytoplasmic TDP-43 De-mixing Independent of Stress Granules Drives Inhibition of Nuclear Import, Loss of Nuclear TDP-43, and Cell Death. Neuron 102, 339–357.e7 (2019).

11. Kim, C. K. et al. Alzheimer’s Disease: Key Insights from Two Decades of Clinical Trial Failures. J. Alzheimers Dis. JAD 87, 83–100 (2022).

12. Aggad, D., Vérièpe, J., Tauffenberger, A. & Parker, J. A. TDP-43 toxicity proceeds via calcium dysregulation and necrosis in aging Caenorhabditis elegans motor neurons. J. Neurosci. Off. J. Soc. Neurosci. 34, 12093–12103 (2014).

13. Buratti, E. & Baralle, F. E. Multiple roles of TDP-43 in gene expression, splicing regulation, and human disease. Front. Biosci. J. Virtual Libr. 13, 867–878 (2008).

14. Yu, S. F. et al. Protein Assembly Modulation: A New Approach to Amyotrophic Lateral Sclerosis (ALS) Therapeutics. *J*. Exp. Neurol. 5, 210–230 (2024).

15. Michon, M. et al. A pan-respiratory antiviral chemotype targeting a transient host multi-protein complex. Open Biol. 14, 230363 (2024).

16. Lingappa, A. F. et al. Small molecule protein assembly modulators with pan-cancer therapeutic efficacy. Open Biol. 14, 240210 (2024).

17. Bechtel, W. & Bich, L. Situating homeostasis in organisms: maintaining organization through time. J. Physiol. 602, 6003–6020 (2024).

18. Tomkins, G. M. The metabolic code. Science 189, 760–763 (1975).

19. Marty, L. et al. The NADPH-dependent thioredoxin system constitutes a functional backup for cytosolic glutathione reductase in Arabidopsis. Proc. Natl. Acad. Sci. U. S. A. 106, 9109–9114 (2009).

20. Noble, D. From Genes to Whole Organs: Connecting Biochemistry to Physiology. in Novartis Foundation Symposia (eds. Bock, G. R. & Goode, J. A.) vol. 239 111–128 (Wiley, 2001).

21. Pardee, A. B. Regulatory Molecular Biology. Cell Cycle 5, 846–852 (2006).

22. Pardee, A. B. & Reddy, G. P.-V. Beginnings of feedback inhibition, allostery, and multi-protein complexes. Gene 321, 17–23 (2003).

23. Fenton, A. W. Allostery: an illustrated definition for the ‘second secret of life’. Trends Biochem. Sci. 33, 420–425 (2008).

24. Müller-Schiffmann, A. et al. A Pan-Respiratory Antiviral Chemotype Targeting a Transient Host Multiprotein Complex. http://biorxiv.org/lookup/doi/10.1101/2021.01.17.426875 (2021) doi:10.1101/2021.01.17.426875.

25. Lingappa, A. F. et al. Small Molecule Assembly Modulators with Pan-Cancer Therapeutic Efficacy. bioRxiv 2022.09.28.509937 (2022) doi:10.1101/2022.09.28.509937.

26. Nardo, G. et al. Amyotrophic lateral sclerosis multiprotein biomarkers in peripheral blood mononuclear cells. PloS One 6, e25545 (2011).

27. Luotti, S. et al. Diagnostic and prognostic values of PBMC proteins in amyotrophic lateral sclerosis. Neurobiol. Dis. 139, 104815 (2020).

28. Pongrácová, E., Buratti, E. & Romano, M. Prion-like Spreading of Disease in TDP-43 Proteinopathies. Brain Sci. 14, 1132 (2024).

29. Duyckaerts, C., Clavaguera, F. & Potier, M.-C. The prion-like propagation hypothesis in Alzheimer’s and Parkinson’s disease. Curr. Opin. Neurol. 32, 266–271 (2019).

30. Wenzhi, Y., Xiangyi, L. & Dongsheng, F. The prion-like effect and prion-like protein targeting strategy in amyotrophic lateral sclerosis. Heliyon 10, e34963 (2024).

31. Feng Yu, S., et al. Protein Assembly Modulation: A New Approach to ALS Therapeutics. http://biorxiv.org/lookup/doi/10.1101/2023.07.24.550252 (2023) doi:10.1101/2023.07.24.550252.

32. Dervishi, I. et al. Protein-protein interactions reveal key canonical pathways, upstream regulators, interactome domains, and novel targets in ALS. Sci. Rep. 8, 14732 (2018).

33. Kanehisa, M., Furumichi, M., Sato, Y., Kawashima, M. & Ishiguro-Watanabe, M. KEGG for taxonomy-based analysis of pathways and genomes. Nucleic Acids Res. 51, D587–D592 (2023).

34. Copley, S. D. Moonlighting is mainstream: paradigm adjustment required. BioEssays News Rev. Mol. Cell. Dev. Biol. 34, 578–588 (2012).

35. Jeffery, C. J. Multitalented actors inside and outside the cell: recent discoveries add to the number of moonlighting proteins. Biochem. Soc. Trans. 47, 1941–1948 (2019).

36. Jia, Y. et al. A Modified Ficoll-Paque Gradient Method for Isolating Mononuclear Cells from the Peripheral and Umbilical Cord Blood of Humans for Biobanks and Clinical Laboratories. Biopreservation Biobanking 16, 82–91 (2018).

