## Supplementary figures and images for "An ALS Assembly Modulator Signature in Peripheral Blood Mononuclear Cells: Implications for ALS Pathophysiology, Therapeutics, and Diagnostics"

### Supplemental Figure 3

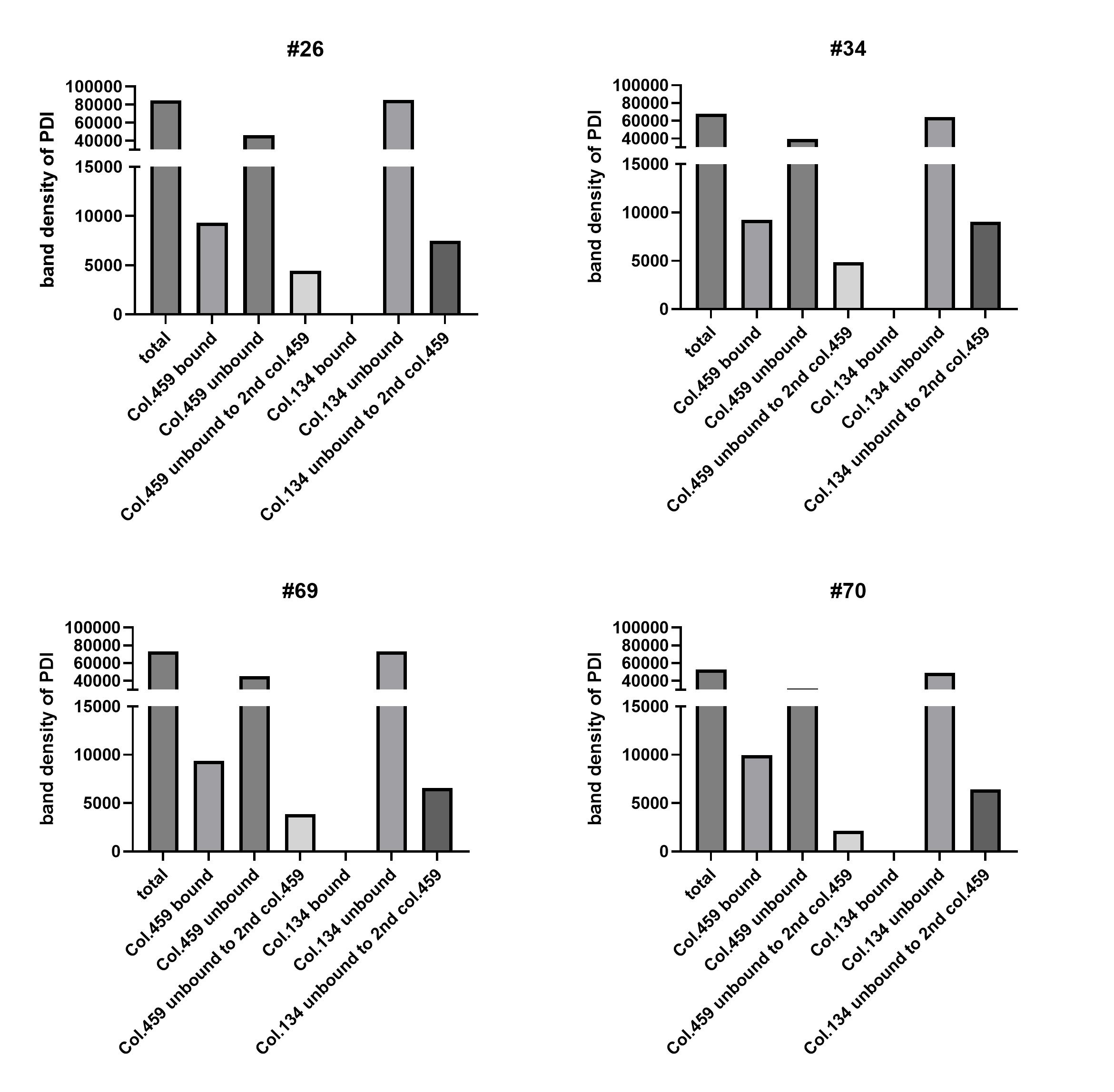
